# Presynaptic vesicles supply membrane for axonal bouton enlargement during LTP

**DOI:** 10.1101/2025.04.29.651313

**Authors:** LM Kirk, GC Garcia, DC Hanka, K Zatyko, T Bartol, TJ Sejnowski, KM Harris

**Affiliations:** Center for Learning and Memory, Department of Neuroscience, University of Texas at Austin, Austin, TX 78712; Computational Neurobiology Laboratory, The Salk Institute for Biological Sciences, La Jolla, CA 92037; Division of Biological Sciences, University of California, San Diego, La Jolla, CA 92093

## Abstract

Long-term potentiation (LTP) induces presynaptic bouton enlargement and a reduction in the number of synaptic vesicles. To understand the relationship between these events, we performed 3D analysis of serial section electron micrographs in rat hippocampal area CA1, 2 hours after LTP induction. We observed a high vesicle packing density in control boutons, contrasting with a lower density in most LTP boutons. Notably, the summed membrane area of the vesicles lost in low-density LTP boutons is comparable to the surface membrane required for the observed bouton enlargement when compared to high-density control boutons. These novel findings suggest that presynaptic vesicle density provides a new structural indicator of LTP that supports a local mechanism of bouton enlargement.

## Introduction

For animals to learn and remember new information, synapses must remain in a modifiable, plastic state. A widely recognized cellular basis for learning and memory is long-term potentiation (LTP), a type of synaptic plasticity that refers to the sustained strengthening of synaptic connections that occurs after brief high frequency stimulation^1^. Synapses are strengthened postsynaptically through insertion of new glutamate receptors into the postsynaptic membrane,^2,3^ while presynaptically there is an increase in the release probability (P_r_)^4,5^. While these changes occur rapidly after LTP induction, they are sustained for hours and are accompanied by structural changes^6,7^.

By two hours, dendritic spines and their excitatory synapses are enlarged at the expense of smaller neighboring spines^8^. Presynaptic boutons, the sites of neurotransmitter release, also undergo structural modifications during LTP^5^. Super resolution imaging of CA3 axons in organotypic slices revealed that boutons were enlarged shortly following LTP induction by high frequency stimulation and remained enlarged for the duration of the experiment^9^. Two hours following LTP induction in acute hippocampal slices, there is a decrease in total bouton number compared to control stimulation, which occurs primarily by losing single synaptic and non-synaptic boutons^10^. In the remaining LTP boutons, both docked and non-docked vesicle counts were decreased at 2 hours compared to control boutons^10,11^. Considering synaptic vesicles are the source of neurotransmitter release, it is perhaps surprising that an increase in P_r_ is accompanied by a decrease in synaptic vesicles. However, during LTP, more of the vesicles at the active zone are tightly docked^7^, likely representing vesicles that are primed and in a readily releasable state^4^. Regardless, the destination of the lost vesicles during LTP remained unclear. Although small transport packets of synaptic vesicles can translocate between neighboring boutons as part of a larger “super pool,”^12^ transport packets are decreased 2 hours following LTP,^11^ and potentiation does not require recruitment of extrasynaptic vesicles^13^. Here, we propose that the membrane from lost synaptic vesicles provides the required increase in surface area for the enlarged presynaptic boutons during LTP. To test this hypothesis, we developed new tools that obtain accurate quantification of object surface areas through 3D reconstruction from serial section electron microscopy (3DEM).

## Results

### Accurate segmentation and quantification of axon surface areas

Accurate measurement of membrane surface area through serial section electron microscopy required a new approach to account for the overlap between obliquely sectioned objects. Membranes of adjacent processes are thin enough (~15 nm) to overlap within the same ultrathin section (~50 nm) when cut obliquely and thus produce an ambiguous grey surface. An *in silico* ultramicrotome was developed to determine how to assign ambiguous membranes to adjacent structures (Figure 1). Overlapping objects from densely segmented neuropil^14^ were re-sectioned virtually to produce membrane boundaries with known surface areas (Figure 1a-d). Accurate surface areas are achieved by assigning the grey surface area to the axon with overlapping cytoplasm on the adjacent sections (Figure 1d, e). Although the membrane surface area of perfectly cross-sectioned boutons can be readily calculated by measuring the perimeter on each section and multiplying by section thickness (Figure 1f-g), most axons are not perfectly cross-sectioned (Figure 1h-i). Thus, the new method allowed an unbiased inclusion of both cross-sectioned and obliquely-sectioned axons to determine the impact of LTP on the membrane surface area of presynaptic boutons.

**Figure 1.**
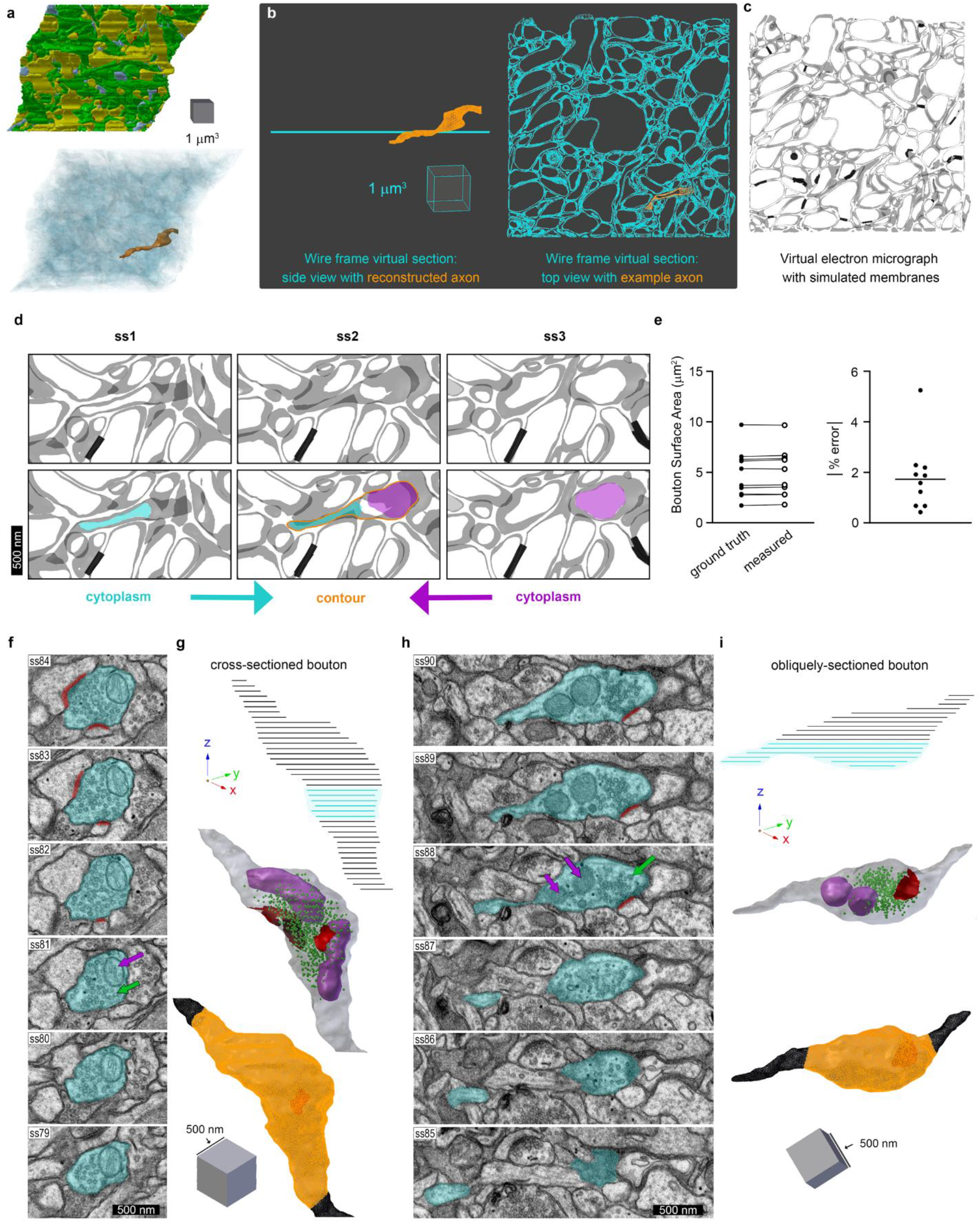
Accurate segmentation and quantification of axon surface areas. **a)** Reconstructed volume containing fully segmented objects from serial section EM (top). Surface meshes were generated using NeuropilTools. A sample axon is shown in orange with the remaining objects in transparent teal (bottom). **b)** Side view and top-down view of one virtual section that was created using the *in silico* ultramicrotome. **c)** Virtual electron micrograph with simulated membranes for each object in (b). **d)** Segmentation strategy using cytoplasm from adjacent serial sections (ss1 and ss3) to guide the amount of grey wall to include in the contour on ss2. **e)** This segmentation strategy resulted in no significant difference between the ground truth values and the manually measured surface area of axons in the virtual volumes (median absolute error = 1.7%, n= 10 axonal boutons). Serial section EM images and 3D reconstructions show example segmentation (teal) of cross-sectioned **(f, g)** and obliquely sectioned **(h, i)** axons, demonstrating how the new segmentation strategy was applied to obtain accurate surface areas. Excitatory postsynaptic densities (red), mitochondria (purple arrows), and synaptic vesicles (green arrows) are also indicated in EM images. **(g, i)** Top: axon contours are shown traversing through the z-axis, and the teal lines indicate the contours shown in the EM example images in **(f)** and **(h)**. Middle: 3D meshes of the axons are generated and smoothed using NeuropilTools. Axons contain synapses (red), presynaptic vesicles (green), and mitochondria (purple). Bottom: the region of each axon used to calculate bouton surface area is indicated in orange, and inter-bouton regions are black.

### Vesicles are lost as boutons enlarge during LTP

Acute hippocampal slices were recovered for at least 3 hours to allow ultrastructure to return to its *in vivo* appearance^15^. Two stimulating electrodes and one recording electrode were placed in the middle of CA1 stratum radiatum (Figure 2a). LTP was induced at one of the electrodes using theta-burst stimulation, and the other electrode received test pulses only as a control^8^ (Figure 2b). In separate slices, LTP was blocked with bath application of 50 µM APV, an NMDA receptor antagonist (Figure 2c). At 2 hours, the slices were rapidly fixed in aldehydes, and tissue near the stimulating electrodes was harvested and processed for serial section EM^8^.

**Figure 2.**
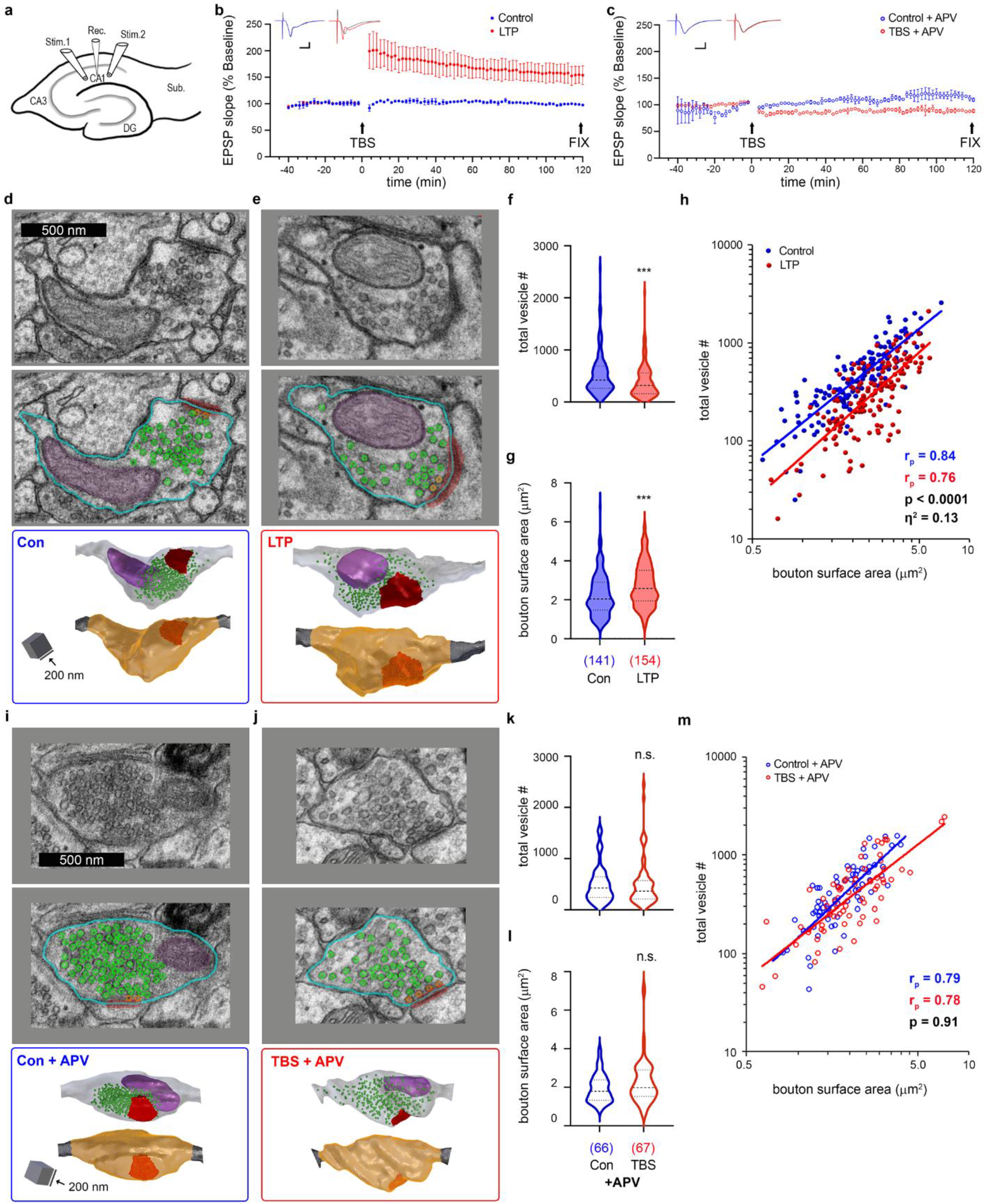
Presynaptic vesicle numbers are decreased, and boutons are larger following LTP. **a)** Diagram of acute hippocampal slice with one recording electrode and two stimulating electrodes delivering alternating stimuli. **b)** EPSP slope increased after theta burst stimulation (TBS) delivered at t = 0 min and was maintained for 2 hours (LTP: red). Test-pulse stimulation does not increase the EPSP (Control: blue). The inset shows average waveforms over 10 minutes from baseline (black; t = −10-0 min) and after TBS or control stimulation (red/blue; t = 110-120 min). **c)** Bath application of 50 µM APV prevents LTP induction with TBS stimulation. For **(b-c)** n = 2 animals per condition, and error bars show SEM. Inset scale bars: x = 5 ms and y = 10 mV. **d-e)** Electron micrographs, segmentations, and 3D reconstructions representing the median vesicle numbers and bouton surface areas for control **(d)** and LTP **(e)**. In addition to axon segmentation (teal), postsynaptic density (red) and mitochondria (purple) segmentations are shown. Docked (orange) and non-docked (green) vesicles are also identified. **f)** Total vesicle number per bouton is decreased with LTP (p = 0.0005). **g)** Bouton surface areas are increased with LTP (p = 0.0001). **h)** Presynaptic vesicle numbers are positively correlated with bouton surface area in both control and LTP conditions (r_p_= Pearson’s R, p<0.0001). Boutons in the LTP condition have fewer presynaptic vesicles compared to control with an effect size (partial eta-squared) of 13% attributed to condition (ANCOVA: F_condition_= 42; df = 1; p<0.0001). For **(f-h)** n_con_=141 and n_LTP_ = 154. **i-j)** Electron micrographs, segmentations, and 3D reconstructions representing the median vesicle numbers and bouton surface areas for control **(i)** and TBS **(j)** in the presence of 50 µM APV. **k-l)** When treated with APV, neither vesicle numbers (p = 0.67) **(k)** nor bouton surface areas (p = 0.12) **(l)** were significantly altered with TBS. **m)** Presynaptic vesicle numbers are positively correlated with bouton surface area in both control and TBS conditions in the presence of APV (Pearson’s R, p<0.0001). However, there was no difference in presynaptic vesicles across bouton sizes between conditions (ANCOVA: F_condition_= .012; df = 1; p = 0.91). For **(k-m)** n_Con+APV_ = 66 and n_TBS+APV_ = 67.

Small synaptic vesicles were located and counted, the synapses and presynaptic mitochondria were segmented, and axonal boutons were delineated along reconstructed excitatory axons (Figure 2d-e, i-j, and extended data Figures 1 and 2). Total vesicle numbers per bouton were decreased with LTP (Figure 2f), and bouton surface areas were increased (Figure 2g). Vesicle numbers and bouton surface areas were positively correlated in both control and LTP conditions, and the drop in vesicle numbers occurred across all bouton sizes after LTP (Figure 2h). When LTP was blocked with APV, there were no differences in vesicle numbers (Figure 2k) or bouton surface areas (Figure 2l) between control and TBS conditions. While vesicle numbers and bouton surface areas were still positively correlated, the relationship between vesicle numbers and bouton surface areas did not differ across the APV control and TBS conditions (Figure 2m). These results indicate that the decrease in presynaptic vesicle number and concurrent increase in bouton surface area require NMDA receptor activation and LTP.

### Decreased vesicle density as a novel structural indicator of LTP

In many of the LTP boutons, non-docked presynaptic vesicles appeared to be spaced farther apart than in the control boutons (Extended Data Figures 1 and 2). To determine whether these spatial relationships were significantly different, a Delaunay mesh was constructed that connected the center points of the vesicles within a bouton (Figure 3a, b). All vesicles could be counted by viewing them through serial sections and distinguishing them from other membrane bound organelles (e.g., smooth endoplasmic reticulum and endosomes). However, some vesicle circumferences could not be measured if their bounding membranes were incomplete or their shape was not spherical. Thus, only the spherical presynaptic vesicles that had an evenly stained membrane with a clear lumen were included to calculate the median vesicle size (see methods; Supplemental Figure 2). The median vesicle radius for each series was then used to calculate the membrane-to-membrane distances from each vesicle to all neighboring vesicles (d_1_, d_2_, d_3_… d_n_) for each non-docked vesicle (v) in the bouton (Figure 3c). The average distance to neighboring vesicles, or DNV, was then determined for each non-docked vesicle (v) within a bouton:

**Figure 3.**
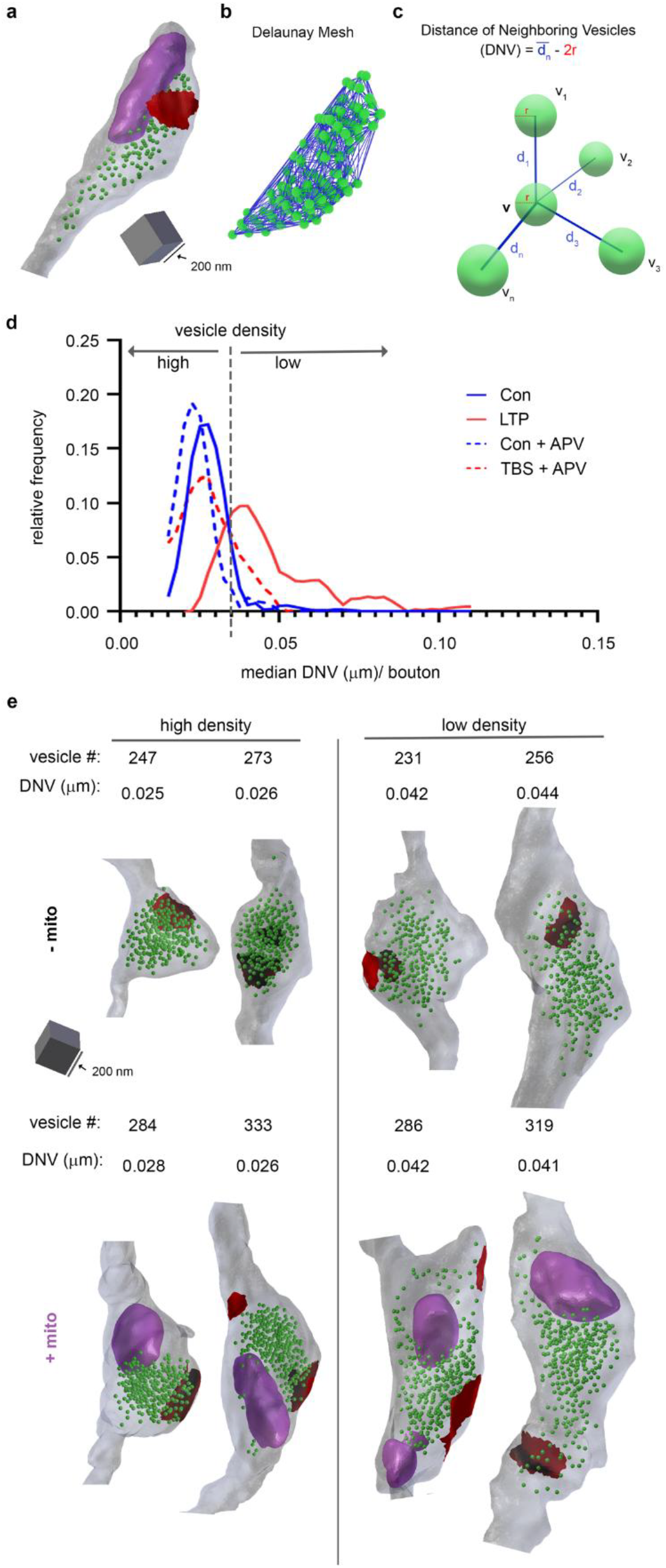
Decreased vesicle density is a feature of LTP. **a)** Example 3D bouton with green vesicles, purple mitochondria, and red synapse. **b)** Delaunay mesh connects the center-points of vesicles in **(a)** to determine the distances of the neighboring vesicles. **c)** Diagram showing how the average membrane to membrane distance to neighboring vesicles (DNV) was calculated for each vesicle. **d)** The relative frequency of the median DNV per bouton is plotted for each condition. The dotted line at 0.035 µm indicates where the LTP and control lines intersect. Boutons with a median DNV ≥ 0.035 µm are classified as having a low vesicle density, and boutons with DNV < 0.035 µm have a high vesicle density. Most boutons in the LTP condition had low density vesicles (77%), compared to the relatively few boutons in control, control-APV, and TBS-APV that had low density vesicles (9%, 6%, 19%, respectively). **e)** Example boutons with low and high vesicle densities that have similar vesicle numbers are shown, indicating that vesicle density is not merely a function of absolute vesicle number. Boutons can have high or low vesicle densities regardless of the presence of mitochondria.

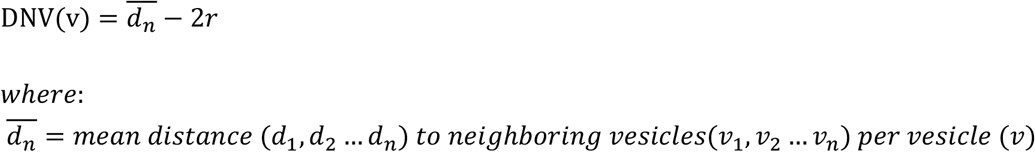

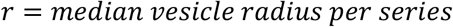

The median DNV was found for each bouton and their relative frequencies plotted for each condition (Figure 3d). Then the value at which the control and LTP curves intersected (0.035 µm) was set as the boundary between boutons with high and low vesicle densities (Figure 3d). In line with our visual observations, there were more LTP boutons with low vesicle densities than in any other condition. Boutons containing comparable vesicle numbers illustrate that the observed high or low vesicle density is not solely determined by the absolute vesicle number (Figure 3e). Mitochondria containing boutons generally have more vesicles (Supplemental Figure 1). However, the presence of a mitochondrion did not affect the proportion of boutons with high or low vesicle densities (Figure 3e, Supplemental Figure 3a). The proportion of single synaptic and multisynaptic boutons that had high and low vesicle densities also did not differ between the LTP and control conditions (Supplemental Figure 3b). Hence, subsequent analyses combined results from all types of presynaptic boutons with and without mitochondria.

### Vesicle Membrane Contribution to Presynaptic Bouton Growth in LTP

We hypothesize that the membrane from the lost synaptic vesicles is used to grow presynaptic boutons during late-phase LTP (Figure 4a). To test this hypothesis, the summed vesicle surface area per bouton was plotted versus the bouton surface area, and the control regression was subtracted from the LTP regression to find the lost vesicle surface area (SA) (Figure 4b):

**Figure 4.**
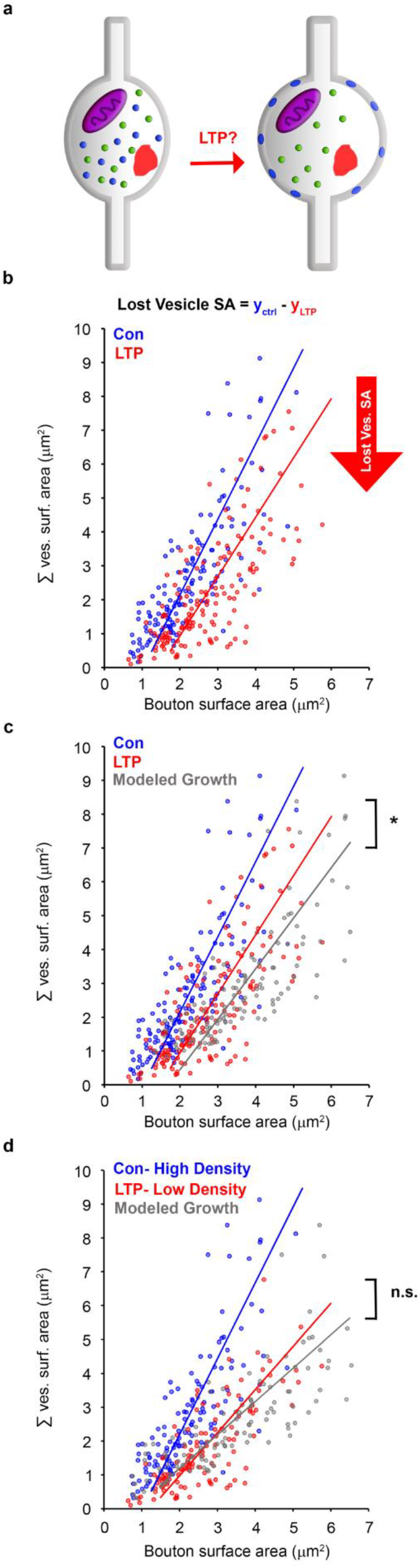
Modeling that includes vesicle density parameters accurately predicts bouton growth from the membrane of presynaptic vesicles lost during LTP. **a)** Illustration demonstrating hypothesis that presynaptic vesicles lost during LTP (blue) provide the necessary membrane to grow presynaptic boutons (right). **b)** Summed vesicle surface area plotted against bouton surface area with model II major axis linear regressions are shown for control and LTP. The equation for lost vesicle surface area with LTP was derived by subtracting the LTP regression from the control regression. **c)** Modeled bouton growth during LTP was determined for each control bouton by using the equation derived in (b) to calculate the predicted summed vesicle surface area lost with LTP and adding it to the control bouton surface area. Modeled boutons and model II linear regression are shown in grey. Although the slopes for the modeled growth and LTP regressions do not differ (p_slope_= 0.69), the intercepts are not equal (p_y-intercept_= 0.4), indicating a poor model. **d)** Control and LTP boutons were restricted to those that had high and low vesicle densities (respectively) as determined in Figure 3. When bouton growth was modeled as in (b-c) using only these restricted data, neither the regression slope nor the y-intercept of the modeled growth boutons differed from the LTP regression (p_slope_= 0.12; p_y-intercept_= 0.20).

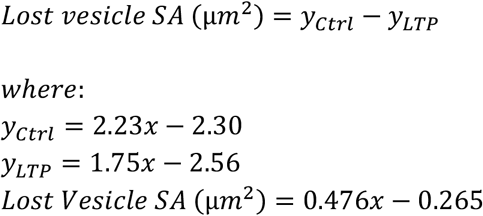

The lost vesicle SA equation was derived to predict the summed vesicle surface area that would be lost for every control bouton in our data set if it were to undergo LTP. To model bouton growth, the calculated lost vesicle surface area was then added to each control bouton surface area. Because we are only measuring unidirectional and unrestricted bouton growth and we know there is a biological ceiling to bouton size, we only plotted modeled boutons that fell within our measured range (Figure 4c, modeled growth). While the membrane from the lost vesicles was sufficient to account for the observed bouton growth during LTP, the model predicted more growth on average than was found experimentally. This discrepancy could be explained by either incomplete incorporation of lost vesicles into the growing boutons or the possibility that some boutons within the LTP condition did not undergo vesicle loss and subsequent growth. To test the latter possibility, the data in our model were restricted to include only control boutons that had high density vesicles (DNV < 0.035 µm) and LTP boutons that had low density vesicles (DNV ≥ 0.035 µm) in accordance with the findings in Figure 3. These density-restricted bouton and summed vesicle surface areas were then plotted, and a new equation for lost vesicle surface area was derived from the regressions:

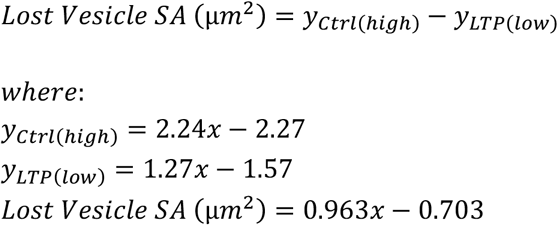

Adding the lost vesicle surface areas to the high-density control boutons resulted in a modeled growth regression that was statistically indistinguishable from the LTP low-density regression (Figure 4d). Therefore, when vesicle density is used as an ultrastructural measure of potentiated boutons, our model supports the hypothesis that lost vesicles are building a bigger bouton during LTP. These novel findings suggest that decreased presynaptic vesicle density is a new structural identifier of potentiated boutons and supports a local mechanism for bouton enlargement.

## Discussion

The current study uses three-dimensional reconstruction from serial section electron microscopy to demonstrate sustained bouton enlargement two hours following theta-burst stimulation, while also replicating prior findings of decreased vesicle numbers ^10,11^. Earlier super-resolution light microscopy studies reported bouton enlargement one hour after high frequency stimulation, an alternative LTP induction paradigm^9^. These combined outcomes suggest that bouton enlargement is a robust form of structural plasticity across different induction protocols and imaging modalities. A key finding of this study is the link between these two phenomena: the boutons exhibiting fewer vesicles post-LTP are also the larger boutons. Moreover, LTP induction leads to decreased vesicle density in a subset of boutons. Notably, these LTP low-density boutons display an increase in surface area that is commensurate with the surface area of lost synaptic vesicles. This novel finding suggests a coordinated structural and functional change.

Individual hippocampal mammalian synapses exhibit a wide variation in the number of small synaptic vesicles, with a significant surplus beyond what is immediately required for synaptic transmission^16^. All vesicles that participate in evoked release comprise the total recycling pool, which are functionally categorized into the readily releasable pool and the recycling pool^17,18^. Synaptic vesicles that are resistant to release even during stimulation are referred to as the resting pool^17,18^. Upon stimulation, vesicles in the readily releasable pool fuse with the plasma membrane, followed by the continued release of vesicles from the recycling pool after the readily releasable pool is depleted. Under baseline conditions, the resting pool does not participate in synaptic transmission. However, within 30 minutes of a potentiating stimuli, resting pool vesicles are recruited into the recycling pool^13,19^. This recruitment process requires both NMDA receptor activation and nitric oxide signaling, indicating the necessity of postsynaptic activity and retrograde communication^13^.

Within the resting pool, vesicles are interconnected by short filaments approximately 30 nm in length^20,21^. Synapsin I, a protein known to form similar 30 nm filaments, links synaptic vesicles to each other and to actin filaments^22^. In Synapsin null *Drosophila* larvae, 30% more vesicles become mobilized, indicating Synapsin as a potential marker for the resting pool of vesicles^23^. Constitutive phosphorylation of Synapsin I by Cyclin-Dependent Kinase 5 (CDK5) enhances its binding to actin, effectively sequestering synaptic vesicles in the resting pool^24^. Conversely, activity-dependent dephosphorylation of Synapsin I by Calcineurin and the subsequent release from actin filaments suggest a mechanism for mobilizing vesicles from the resting pool to the recycling pool upon neural activity^13,24^. In this study, we identified vesicle density as a novel presynaptic structural marker of potentiation. Interestingly, the threshold used to categorize boutons as having high or low vesicle density was based on a distance to neighboring vesicles (DNV) of 35 nm, a value notably comparable to the length of Synapsin filaments. It is tempting to speculate that vesicles in the low-density boutons were mobilized by the activity-dependent dephosphorylation of Synapsin 1, recruiting vesicles from the resting pool to grow the bouton. While previous studies focused on recruitment of vesicles at earlier timepoints with LTP (≤ 30 minutes), this is the first evidence that similar dynamics may be at play during late-phase LTP (2 hours).

To prevent the depletion of the vesicle pool, which would halt neurotransmission, exocytosis is tightly coupled with endocytosis. Following vesicle release at mammalian synapses, membrane is rapidly retrieved to maintain the vesicle pool through two main mechanisms^18^. The first mechanism, ultrafast endocytosis, is a quick, localized process occurring on the millisecond timescale^25^. The second mechanism, activity-dependent bulk endocytosis (ADBE), retrieves larger amounts of membrane over a few seconds^26^. Interestingly, ADBE requires activity-dependent dephosphorylation of Dynamin I by Calcineurin^27^, suggesting a shared mechanism with the previously described role of Calcineurin in mobilizing resting pool vesicles through Synapsin I dephosphorylation. If vesicle membrane is immediately retrieved after intense stimulation, then how does vesicle membrane remain within the plasma membrane to enlarge boutons with LTP? These results initially seem contradictory with the findings presented here. However, there are some key distinctions that could explain these phenomena. First, ADBE operates on a completely different timescale than late-phase LTP (seconds vs. hours). This significant temporal difference suggests that ADBE, operating in the immediate aftermath of intense stimulation, likely serves a different homeostatic role than the longer-lasting structural changes observed during late-phase LTP. Second, the TBS protocol used to induce LTP in this study may engage different signaling pathways than the prolonged bursts of stimulation used to elicit ADBE^26^. Brain-derived neurotrophic factor (BDNF) is a well-known player in late-phase LTP, and its secretion is sensitive to alterations in stimulation paradigms, with LTP-producing stimuli eliciting increased BDNF release^28,29^. Interestingly, BDNF has been shown to prevent the re-phosphorylation of Dynamin I, thereby inhibiting the rapid membrane retrieval mechanisms of ADBE^30^. This BDNF-mediated inhibition of ADBE represents one potential mechanism by which membrane retrieval might be modulated during longer-lasting plasticity, allowing for the incorporation of membrane necessary for the observed bouton enlargement. To our knowledge, there have not been any studies directly examining the mechanisms of endocytosis during LTP-inducing stimuli, whether it be TBS or HFS, leaving this an open area for future investigation.

Given the observed sustained bouton enlargement following LTP induction, it is important to understand its functional implications. Early modeling work suggested that alterations in axon geometries can influence action potential propagation by affecting its amplitude and width^31^. More recently, modeling has demonstrated that increased axon diameter accelerates action potential conduction velocity between boutons; conversely, larger bouton size decelerates it^9^. Furthermore, activity-dependent plasticity has been shown to modulate action potential latency and amplitude hours after induction^32^. Considering the importance of precise action potential timing for spike-timing dependent plasticity, changes in bouton size could represent a key mechanism for fine-tuning the arrival of these signals at postsynaptic sites. Beyond conduction velocity, bouton morphology also influences local calcium dynamics within the bouton and adjacent inter-bouton regions^33^, thereby having an impact on vesicle fusion and local signaling cascades that occur during plasticity. Thus, the LTP-induced bouton enlargement we observed likely contributes to both the temporal and biochemical modulation of synaptic transmission.

One remaining question is how bouton growth and decreased vesicle density relates to postsynaptic spine and synapse enlargement with LTP^8^. Previous work showed that boutons with smaller synapses and vesicle pools display greater mobilization of vesicles from the resting pool to the recycling pool compared to boutons with larger synapses and vesicle pools^13,19^. Since the low-density boutons observed in this study likely correspond to boutons with greater vesicle mobilization, low-density boutons are unlikely to form synapses preferentially with potentiated spines that have larger synapses. Whether enlarged boutons contribute to the remodeling of synaptic clusters near larger spines that have reached the final capacity for growth remains an open and compelling area for future investigation^34^. The manual segmentation of vesicles in large-scale 3DEM data sets is tedious and provides a substantial hurdle to a more comprehensive circuit-level analysis needed to address these questions more generally. Although the incorporation of machine learning into 3DEM automatic segmentation pipelines has made many recent advances^35,36^, accurate segmentation of synaptic vesicles remains a challenge. Dysregulation of long-term presynaptic plasticity is emerging as an important player in several neuropsychiatric and neurodegenerative diseases^37,38^. This study reveals new evidence demonstrating the dynamic structural plasticity of presynaptic boutons, with significant implications for both normal brain function and neurological disorders.

## Methods

### Animals

Serial section images were analyzed from series that had been prepared for prior publications as described below. Procedures were carried out in accordance with the National Institutes of Health guidelines for the humane care and use of laboratory animals. All protocols and procedures involving animals were approved by the Institutional Animal Care and Use Committee at The University of Texas at Austin. All 3DEM data were collected from adult male Long-Evans rats.

### Physiology

Slice preparation and physiology were previously described and published^8,11^. Adult male Long-Evans rats aged 60-61 days old were anesthetized with halothane and decapitated, and 400 µm thick slices were rapidly collected from the middle third of the hippocampus at room temperature using a Stoelting tissue chopper. Slices were recovered in a humidified interface chamber using artificial cerebrospinal fluid for approximately 3 hours at 32^°^C. For experiments where LTP was blocked, 50 µM APV (D-2-amino-5-phosphonopentanoic acid; Sigma Aldrich, St. Louis, MO) was added into the interface chambers at the end of the 3 hour recovery period and remained for the duration of the experiment (Figure 2c). One extracellular recording electrode was placed in the middle of stratum radiatum, and two bipolar stimulating electrodes were placed ~300-400 µm on either side of the recording electrode (Figure 2a). An input-output curve was generated, and the initial slope of the field excitatory postsynaptic potential (fEPSP) was measured. The stimulus intensity was set at 75% of the maximum fEPSP, just below the population spike threshold, and held constant throughout the experiment. Stimulations were delivered once every 2 min, with a 30 second lag between control and LTP stimulation electrodes. Baseline fEPSPs were collected for 30 minutes before theta-burst stimulation (TBS: 8 trains spaced at 30 second intervals, with each train consisting of 10 bursts at 5 Hz, and each burst consisting of 4 pulses at 100 Hz) was delivered to one stimulating electrode to produce LTP (Figure 2b-c). The fEPSPs were recorded in response to alternating control and LTP test pulses for 2 hours. The sites of LTP versus control stimulation (near CA3 or subiculum) alternated between experiments.

### Tissue fixation, processing, and imaging

Fixation, processing, and imaging methods were previously described and published^8^. Briefly, electrodes were removed, and slices were rapidly fixed by immersing them into mixed aldehydes (6% glutaraldehyde, 2% paraformaldehyde in 100 mM cacodylate buffer with 2 mM CaCl_2_ and 4 mM MgSO_4_) and microwaving for 10 sec. Slices were then stored in the same fixative overnight at room temperature before being embedded in 7% agarose and then vibra-sliced at 70 µm (Leica VT 1000S, Leica, Nussloch, Germany). Vibra-slices that showed a visible surface indentation from the stimulation electrodes along with 2 adjacent vibra-slices were identified and processed using the following procedure: 1% osmium/1.5% potassium ferrocyanide mixture, 1% osmium alone, dehydrated through graded ethanol (50-100%) and propylene oxide, embedded in LX112, and placed in a 60°C oven for 48 hours.

Test thin sections were cut from regions located 150-200 µm lateral to the stimulating electrodes and 120-150 µm beneath the air surface of the slice, and mounted on Pioloform-coated slot grids (Synaptek, Ted Pella). Sections were counterstained with saturated ethanolic uranyl acetate and Reynolds lead citrate and imaged on a JEOL 1230 electron microscope with a Gatan digital camera at 5,000X magnification. Images were then assessed for quality of tissue preservation. Approximately 200 serial sections were then cut and imaged, and a five-letter code was used to mask the identity of experimental conditions in subsequent analyses. A diffraction grating replica (Ernest Fullam, Latham, NY) was imaged with each series to calibrate X-Y dimensions.

### *In silico* ultramicrotome

The *in silico* ultramicrotome is a software tool designed to assess the accuracy of the reconstruction pipeline starting from hand-tracing of the electron microscopy (EM) images through the final step of mesh generation. It is an open-source Python script built on top of Blender, a popular 3D CAD/CGI platform. Ground truth values for accuracy testing of our reconstruction methods were established by generating surface meshes from densely segmented hippocampal neuropil^14^ in Blender, using NeuropilTools and Contour Tiler, and calculating surface areas for each object (Figure 1a). These objects were re-sectioned using the *in silico* ultramicrotome to create wire-framed virtual sections of known thickness (Figure 1b). Simulated membranes were then added to objects to create virtual electron micrographs (Figure 1c). Objects in the virtual micrographs can then be re-traced (Figure 1d), their meshes regenerated, and their surface areas compared to ground truth values.

### Unbiased axon selection and segmentation

Series were initially aligned, and five dendrites of similar caliber and their synapses were previously segmented using legacy Reconstruct software^8,39^. Section thickness was computed by dividing the diameters of longitudinally sectioned mitochondria by the number of sections they spanned^40^. Series were then imported into PyReconstruct^41^. For each dendrite, the middle 15 excitatory spine synapses were identified, and their axons were traced. Most axons traverse the neuropil in a winding fashion, meaning their membranes are often not cut in cross section and their edges are ambiguous to discern. We used the *in silico* ultramicrotome to test different segmentation methods and found that the most accurate segmentation could be achieved by including grey wall membrane that overlapped with cytoplasm on adjacent sections (Figure 1d-e). Axonal boutons were traced to a sufficient distance to capture both the target bouton and adjacent inter-bouton regions. If the axon made another bouton nearby, then that bouton and its synapses were also traced. Any additional PSDs that contacted the traced boutons were segmented. Both docked and non-docked vesicles were stamped, and mitochondria were traced.

### Bouton measurement in NeuropilTools

Axon traces were imported from Reconstruct into Blender as contours using NeuropilTools in CellBlender (a modeling plug-in for Blender)^42^. The contours were converted into a three-dimensional mesh using NeuropilTools via its Contour Tiler module. The resulting mesh was then improved using the GAMer module to smooth the mesh and remove distortions. Synaptic contact areas were tagged (via the obj_tag_region module) onto the axon surface areas and non-docked synaptic vesicles and mitochondria were imported for visualization. Two experimenters, masked as to condition, examined each axon in 3D to assess morphology from all angles. The lasso tool (included in NeuropilTools) was used to select the bouton region (Figure 1g,i). Bouton ends were defined as the region where constriction ceased and a relatively uniform tubular circumference was maintained along the axon (inter-bouton region). The mitochondria, vesicles, and synapses provided additional context, especially along the slopes between the axonal bouton and inter-bouton regions. Each experimenter measured each bouton twice, and intra- and inter-tracer errors were determined for the surface area. When measurement error was above 10%, the bouton was reviewed by both tracers in tandem to determine the cause of the discrepancy. Boutons exhibiting ambiguous morphology, such as those appearing to merge or split, were excluded from analysis. The average intra-tracer error was 2.04% +/− 0.08, and the average inter-tracer error was 2.43% +/− 0.14. The average of the four surface area measurements was used for analysis.

### Vesicle surface area measurements

Although all synaptic vesicles within the selected boutons were identified, stamped, and counted, only a subset of vesicles were manually segmented to obtain their surface areas. Because we used circumference to calculate the surface area, it was important to only include vesicles that were circular in shape, clearly distinguishable from other vesicles, and were fully contained within a single section (which tend to appear with a distinct membrane and a clear lumen; Supplemental Figure 2a-c). To sample from the range of bouton sizes and conditions for vesicle measurement, axonal boutons from each series were first categorized by the presence of mitochondria. Each population was then ranked by bouton surface area and placed into 10% bins. A bouton was randomly selected from each bin and if it contained a minimum of 10 vesicles that met our criteria, the vesicles were segmented and the circumference used to calculate spherical surface area. If a selected bouton did not contain at least 10 vesicles matching our criteria, another was selected from the same size bin instead. A total of 30 boutons containing mitochondria and 30 boutons lacking mitochondria in each series had their vesicles traced (Supplemental Figure 2d; n=8 series, 480 boutons, 8,629 vesicles).

### Distance of neighboring vesicles

To compute the distance between neighboring vesicles, we used the Delaunay triangulation associated with the vesicle cluster. The triangulation was generated using the associated routine in SciPy 1.1.0 through a Python script written for Blender 2.79. The Euclidean distances between neighboring vesicles (i.e., vesicles connected in the Delaunay triangulation) were calculated and the mean distance was computed for each vesicle. Because the distribution of mean vesicle distances within each cluster was skewed, the median was reported for each bouton. Finally, we obtained a membrane-to-membrane distance by subtracting two times the median vesicle radius for each series (see previous “Vesicle surface area measurements”).

### Modeled bouton growth

The summed vesicle surface area per bouton was calculated by multiplying each bouton’s vesicle number (Figure 2f) by the median vesicle surface area of the originating series (Supplemental Figure 2d). Outliers were defined as values exceeding three times the interquartile range (IQR) beyond the lower and upper quartiles for both summed vesicle surface area and bouton surface area. Using these criteria, three data points were removed from the control condition, and one from the LTP. Bouton surface areas were then plotted against summed vesicle surface areas. We used GraphPad Prism 10 to determine model II Major Axis linear regressions, suitable when both X and Y variables are in the same units and subject to measurement error. For comparison of variable relationships between LTP and Modeled Growth, we first tested for the homogeneity of slopes using the F-test on the interaction term. The slopes were not significantly different, so we proceeded to test for differences in the elevations (y-intercepts) of the lines. This allowed us to determine if the groups differed significantly in their Y values after accounting for the influence of the X variable.

## Statistical analysis

Analysis of covariance (ANCOVA) was used to test for differences in vesicle counts between control and LTP conditions across bouton surface areas (Figure 2 and Supplemental Figure 1). Because both vesicle counts and bouton surface areas have skewed distributions, data were log10 transformed prior to analysis. Statistica (TIBCO Software Inc. (2020), Data Science Workbench, version 14, http://tibco.com) was used to first test for ANCOVA assumptions (Levine’s test for homogeneity of variances, homogeneity of slopes). In Supplemental Figure 1d, Levine’s test showed a slightly significant difference in variance (p = 0.045). However, we had roughly equal sample sizes between conditions, and the variance ratio was less than 4 (1.5), so we determined this to be a minor violation and continued with the ANCOVA analysis. Although ANCOVAs were performed on log transformed data, the raw data points were plotted on log axes with a power regression in Microsoft Excel to visualize the skewed nature of the raw data and the linear relationships.

GraphPad Prism was used for all other statistical analyses (Version 10.3.1 for Window, Boston, Massachusetts USA, www.graphpad.com). In Figure 3d, frequency distributions for the median distance to neighboring vesicles per bouton were determined using 2.5 nm bins. Relative frequency histograms were then created using 2^nd^ order smoothing (4 neighbors). The Mann-Whitney test was used for all pairwise comparisons. For all figures, *p<0.05, **p<0.01, ***p<0.001, ****p<0.0001.

## Author contributions

The work was conceptualized by LMK and KMH. LMK, DCH, and KZ acquired data. Data was analyzed by L.M.K. and G.C.G. LMK interpreted all data and wrote the manuscript with input from all co-authors. T.B. created new software used in the study. T.J.S. and K.M.H. obtained funding and supervised the project.

## Acknowledgements

This study was supported by NIH Grants R01MH095980, R56MH139176, and National Science Foundation NeuroNex Grants 2014862 and 2219894 to K.M.H. We thank Patrick Parker for proofreading and helpful suggestions on the manuscript.

---

**Extended Data Figure 1.**
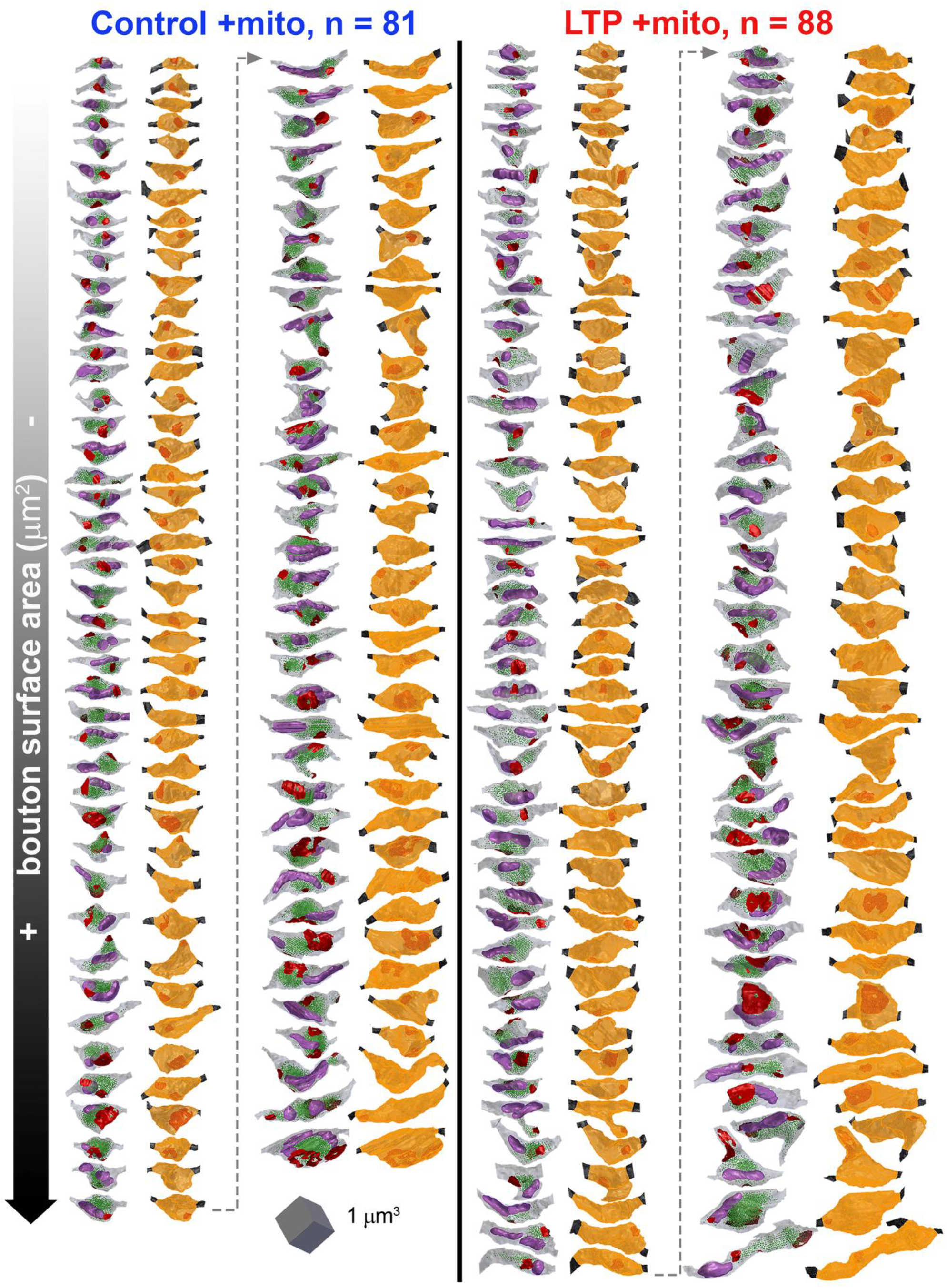
Mitochondria containing boutons. All boutons containing mitochondria that were measured for this study are shown in descending order from smallest to largest surface area for control (left) and LTP (right) conditions. Each bouton is shown as a pair with its 3D rendering containing presynaptic vesicles (green), mitochondria (purple), and synapses (red) on the left and the selected region of the bouton used for quantification on the right (orange).

**Extended Data Figure 2.**
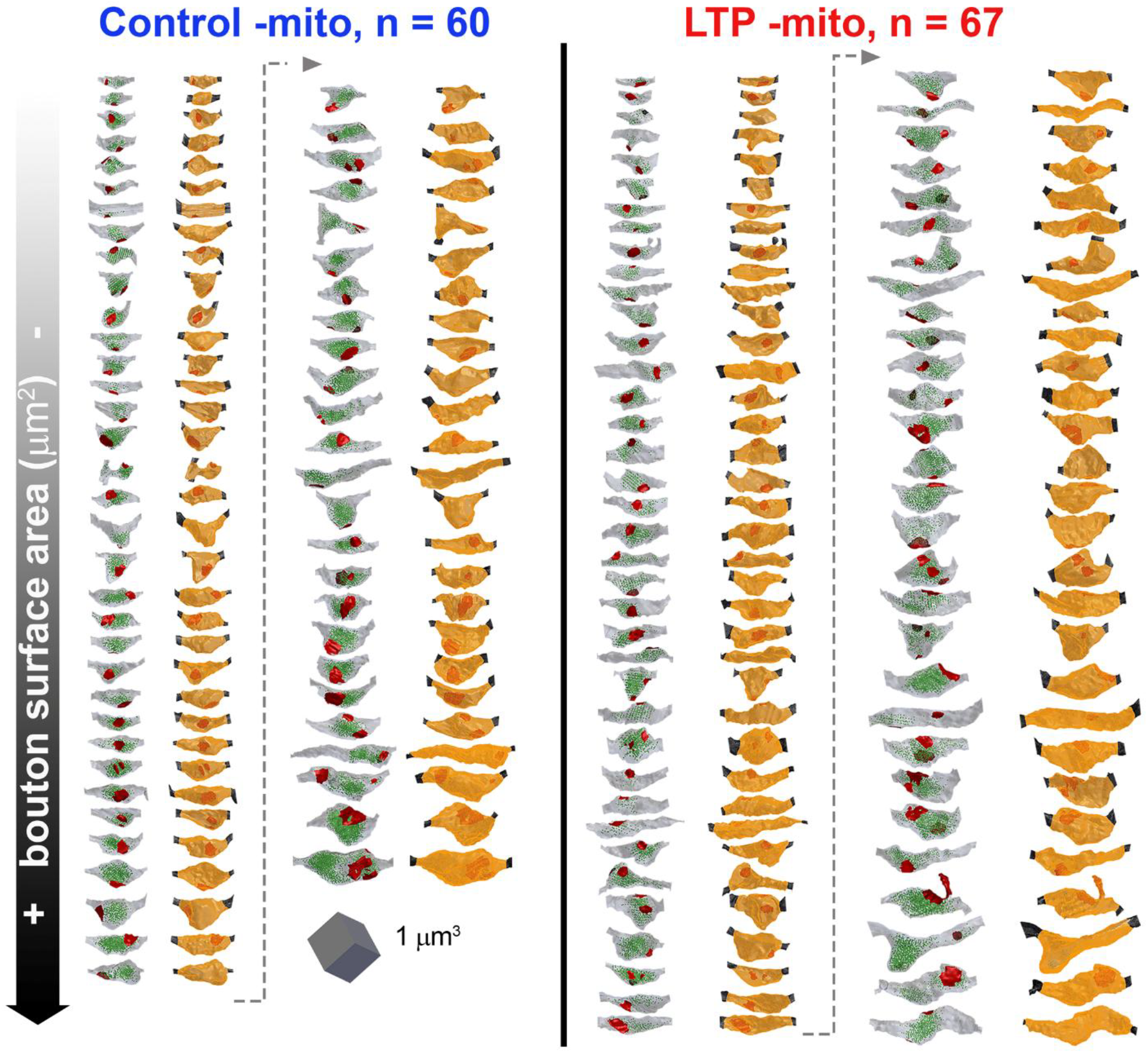
Mitochondria lacking boutons. All boutons lacking mitochondria that were measured for this study are shown in descending order from smallest to largest surface area for control (left) and LTP (right) conditions. Each bouton is shown as a pair with its 3D rendering containing presynaptic vesicles (green) and synapses (red) on the left and the selected region of the bouton used for quantification on the right (orange).

**Supplemental Figure 1.**
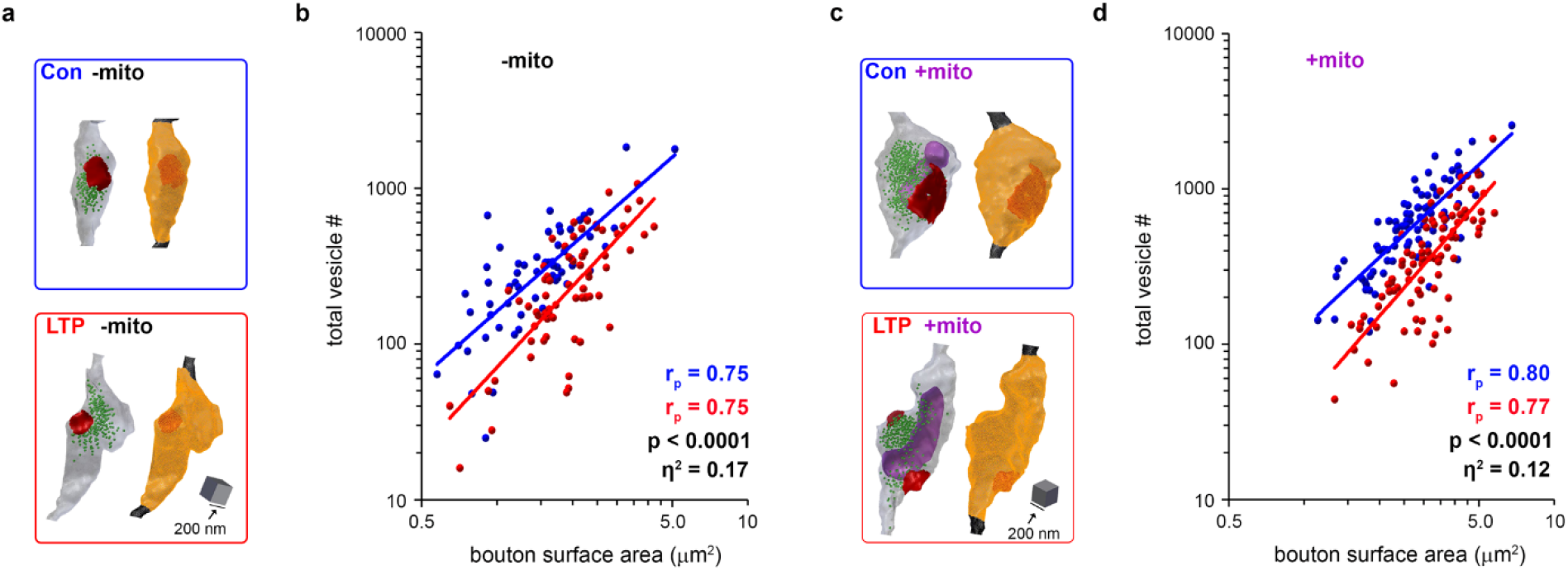
Boutons with and without mitochondria show vesicle loss with LTP. **a)** 3D reconstructions representing the median vesicle number and surface area of control and LTP boutons lacking mitochondria. **b)** In mitochondria lacking boutons, presynaptic vesicle numbers are also positively correlated with bouton surface area in both control and LTP conditions (Pearson’s R, p<0.0001). Similarly, LTP boutons lacking mitochondria have fewer presynaptic vesicles across bouton sizes compared to control, with an effect size of 17% attributed to condition (ANCOVA: Fcondition= 26; df = 1; p<0.0001; nCon = 60; nLTP = 67). **c)** 3D reconstructions representing the median vesicle number and surface area of control and LTP boutons containing mitochondria. **d)** In mitochondria containing boutons, presynaptic vesicle numbers are positively correlated with bouton surface area in both control and LTP conditions (Pearson’s R, p<0.0001). LTP boutons containing mitochondria have fewer presynaptic vesicles across bouton sizes compared to control, with an effect size (partial eta-squared) of 12% attributed to condition (ANCOVA: F_condition_= 23; df = 1; p<0.0001; n_Con_ = 81; n_LTP_ = 87).

**Supplemental Figure 2.**
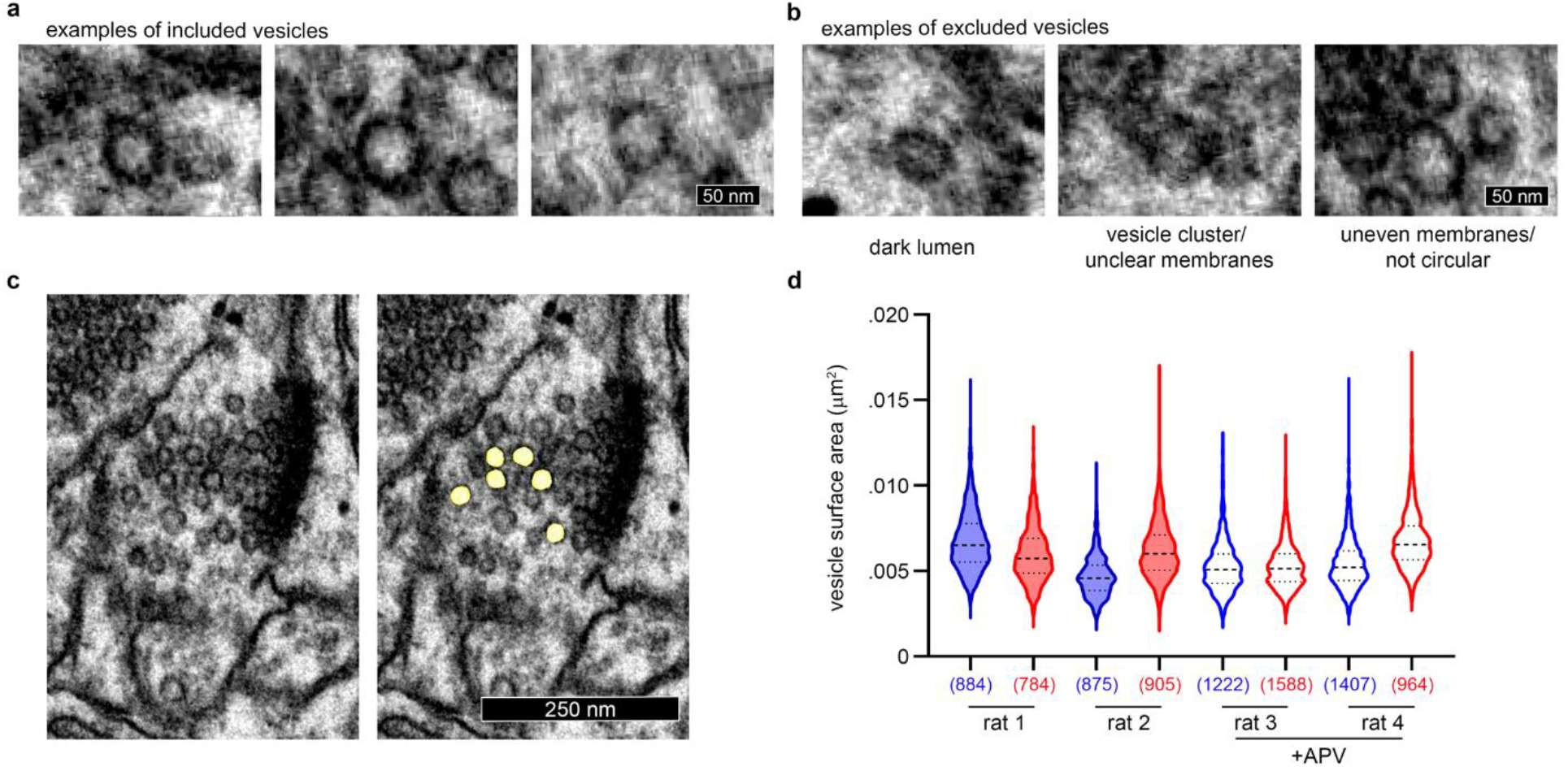
Vesicle inclusion and exclusion criteria for surface area measurements. Examples of presynaptic vesicles that met inclusion criteria for segmentation: clearly distinguishable from other vesicles, circular in shape, and evenly stained membrane with clear lumen. **b)** Examples of synaptic vesicles that did not meet our criteria. Reasons for exclusion are listed below each image. **c)** Example bouton that was selected for vesicle surface area measurement. The vesicles that passed our conservative inclusion criteria and used for quantification are indicated in yellow (right). **d)** Median vesicle surface areas are shown for each series +/− IQR, with (n = number of vesicles) in parenthesis below each group.

**Supplemental Figure 3.**
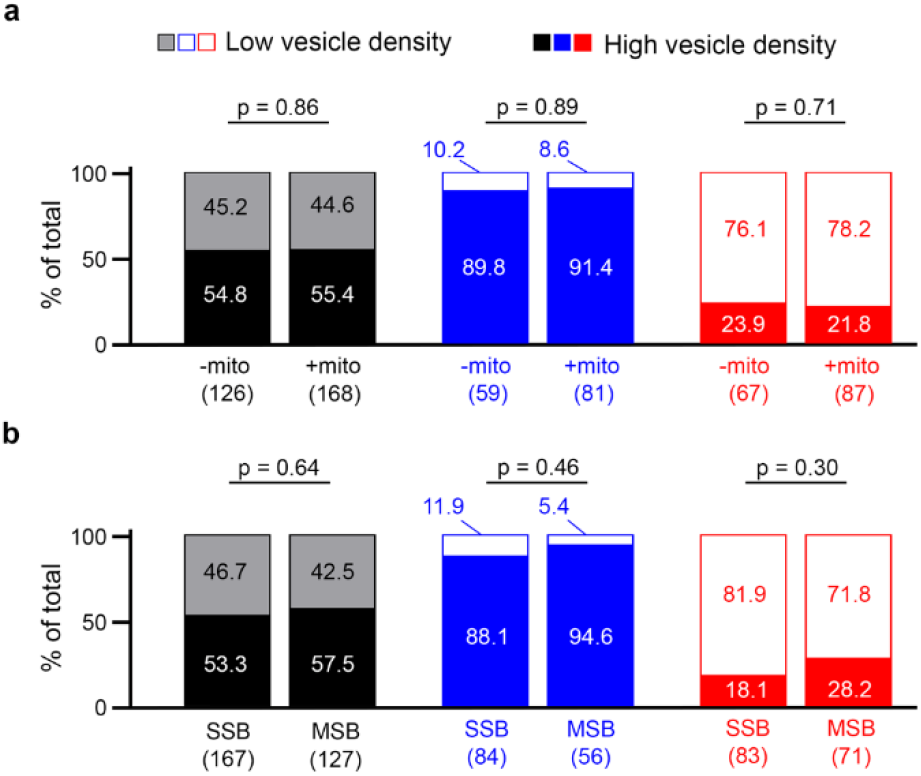
Proportion of boutons with high and low vesicle densities does not depend on mitochondria or number of synapses. **a)** When looking at all boutons in our data set (grey/black, left), the presence of mitochondria did not affect the proportion of boutons that had high or low vesicle densities (chi-squared, p = 0.86). Similarly, there was no difference between −/+ mitochondria boutons in either control (blue, p = 0.89) or LTP (red, p = 0.71) conditions. **b)** Similarly, whether a bouton was single synaptic (SSB) or multi-synaptic (MSB) had no effect on the proportion of high/low vesicle densities in all boutons (p = 0.64), control (p = 0.46), or LTP (p = 0.30) conditions.

